# Atomic protein structure refinement using all-atom graph representations and SE(3)-equivariant graph neural networks

**DOI:** 10.1101/2022.05.06.490934

**Authors:** Tianqi Wu, Jianlin Cheng

**Affiliations:** Department of Electrical Engineering and Computer Science, University of Missouri, Columbia, MO 65211

## Abstract

Three-dimensional (3D) protein structures reveal the fundamental information about protein function. The state-of-art protein structure prediction methods such as Alphafold are being widely used to predict structures of uncharacterized proteins in biomedical research. There is a significant need to further improve the quality and nativeness of the predicted structures to enhance their usability. Current machine learning methods of refining protein structures focus mostly on improving the backbone quality of predicted structures without effectively leveraging and enhancing the conformation of all atoms including side-chain, while molecular simulation methods are computationally intensive and time-consuming.

In this work, we develop ATOMRefine, a deep learning-based, end-to-end, all-atom protein structural model refinement method. It uses a SE(3)-equivariant graph transformer network that is equivariant to the rotation and translation of 3D structures in conjunction with a novel graph representation of all atoms to directly refine protein atomic coordinates of all the atoms in a predicted tertiary structure represented as a molecular graph. The method is first trained and tested on the structural models in AlphafoldDB whose experimental structures are known, and then blindly tested on 69 CASP14 regular targets and 7 CASP14 refinement targets. ATOMRefine improves the quality of both backbone atoms and all-atom conformation of the initial structural models generated by AlphaFold. It also performs better than the state-of-the-art refinement methods in multiple evaluation metrics including an all-atom model quality score – the MolProbity score based on the analysis of all-atom contacts, bond length, atom clashes, torsion angles, and side-chain rotamers. As ATOMRefine can refine a protein structure quickly, it provides a viable, fast solution for improving protein geometry and fixing structural errors of predicted structures through direct coordinate refinement.

## Introduction

Every cell in the human body contains proteins. Protein participates in most cellular processes, ranging from DNA replications to immune responses. Protein functions are intimately connected with their three-dimensional shapes. Therefore, predicting the protein structure from sequence has been a long-standing grand challenge in computational biology. Recently, AlphaFold^1,2^ is shown to predict highly accurate tertiary structures for most proteins, which is considered a big advance in the field. However, there are still some limitations in the AlphaFold predicted structures. The recent application of AlphaFold2^3^ to predicting the structures in the human proteome showed that the conformation of 58% of the total residues was of high accuracy with the predicted confident score pLDDT ^1^> 70, leaving the rest 42% of the total residues with the confidence score pLDDT ≤ 70. Besides, a strong correlation between the Alphafold model quality and the availability of homologous templates in the Protein Data Bank (PDB) has been observed in a few benchmarking studies^4–6^, suggesting that there is still a room to improve the quality of AlphaFold models, particularly for proteins without homologous templates in the PDB. Moreover, current protein structure prediction methods including AlphaFold have been focused on predicting the backbone structure of proteins correctly without emphasizing improving the nativeness and all-atom geometry of predicted structures, leaving significant room to improve the all-atom quality of predicted structures^7^. Therefore, there is a significant need to further refine the protein structures predicted by state-of-the-art methods such as AlphaFold to improve their usability in biomedical research.

Currently, typical model refinement methods apply molecular dynamics (MD) simulation, energy minimization, or fragment assembly to refine input protein structures. Successful MD-based methods^8–11^ are physics-based approaches to sampling multiple MD trajectories following the physical principles regarding atomic interactions, which are computation-intensive and time-consuming. Energy minimization-based methods^7,12^ focus on repacking the backbone and side-chain atoms with composite physics and knowledge-based force fields. Fragment assembly-based methods are like knowledge-based methods, taking advantage of template fragment information in the PDB as well as statistical potentials. A notable method is ROSETTA^13^, which uses predicted estimated local structural errors to inform the fragment assembly, followed by side-chain rebuilding and energy minimization in all-atom representation. Though those methods prove to be effective in the refinement of some protein structures, they require extensive conformation sampling and a lot of computing resources.

Deep learning has recently been applied to improve the geometric property of the protein three-dimensional structure. Graph neural networks were used by GNNRefine^14^ to refine the backbone atoms of protein structure. It largely relies on a Rosetta protocol for the full-atom model reconstruction. In the refinement module of RoseTTAfold^15^, a SE(3)-equivariant graph transformer^16^ is used to refine backbone atoms without directly using machine learning to leverage and improve side-chain atoms in a protein structure. It produces a refined model with only backbone atoms and cannot be used as a standalone tool to refine a third-party model.

Inspired by the application of geometric deep learning to molecular structure prediction that can avoid the expensive and extensive conformation sampling, here we present ATOMRefine, a new SE(3)-equivariant transformer network based on a novel all-atom representation of atom types, amino acid types, atom-atom distances, and covalent bonds for refining protein structures in the full-atom scale. Its graph representation of all the atoms of a protein structure enables the network to leverage sequence-based and spatial information from the entire protein structures to update node and edge features and catch the global and local structural variation from the initial model to the native structure iteratively. The 3D-equivariance makes it possible for ATOMRefine to learn essential structural properties regardless of the rotation and translation of the input structure. The network outputs the refined coordinates of all the atoms directly, without using any external protein full-atom reconstruction protocol. To the best of our knowledge, ATOMRefine is the first end-to-end all-atom 3D-equivariant network approach to refine the protein model prediction on the full atom scale.

Evaluated on both AlphaFold and the 14^th^ Critical Assessment of Techniques for Protein Structure Prediction (CASP14) datasets, ATOMRefine improves the quality of both backbone and all atoms of the initial structural model in terms of GDT-TS score, GDT-HA score, RMSD, and Molprobity. Noticeably, ATOMRefine can maintain or improve the model quality over the initial models generated by AlphaFold and the existing model refinement methods, and generate far fewer model degradation cases than the others.

## Results

### Comparison of ATOMRefine with other refinement methods in terms of backbone quality

Geometric deep learning-based approaches have been applied to protein structure refinement, among which GNNRefine yields some quality improvement from initial models. However, its machine learning component heavily focuses on the backbone atom refinement, and largely relies on the Rosetta refinement protocol for the final full-atom refinement. In contrast, ATOMRefine applies an all-atom SE(3)-equivariant graph transformer to directly refine all the atoms of a protein structure. Directly refining all the atoms has the benefit of generating an all-atom refined structure in an end-to-end fashion, but it requires a much larger molecular graph to represent all the atoms in a protein structure than that representing only backbone atoms (or only Ca atom). To investigate the trade-off of using a full-atom representation, we implement two versions of our method based on the same SE(3)-equivariant graph transformer architecture: (1) ATOMRefine – the all-atom refinement method and (2) ATOMRefine_backbone – the backbone atom refinement method. Both of them are trained and validated on the same dataset.

We evaluate ATOMRefine, ATOMRefine_backbone, GNNRefine, and a widely-used energy minimization-based method – ModRefiner on the AlphaFoldDB test set and the structural models of 69 CASP14 targets. For the Alphafold DB test set, the structural models from the Alphafold DB are used as the initial models. For the CASP14 dataset, Alphafold2 is used to predict the structures of the CASP14 targets that are used as the initial models. For each initial model, the best of five refined models produced by each method is selected for evaluation against the true experimental structures. The backbone quality of the initial models and the models refined by these methods is reported in **Table 1**.

**Table 1.** Performance of ATOMRefine, ATOMRefine_backbone, GNNRefine, and ModRefiner on AlphaFold DB test set and CASP14 dataset in comparison with initial models. Bold numbers denote the best results. Improvement percentage denotes the percentage of the models that have been improved by each method in terms of the GDT-HA score.

| Test set | Method | GDT-TS | GDT-HA | RMSD of $C\alpha$ | Improvement percentage (%) |
| --- | --- | --- | --- | --- | --- |
| AlphaFoldDB test set | Initial model | 81.42 | 69.84 | 4.09 | / |
|  | ATOMRefine | 81.59 | 70.04 | 4.08 | **56.48%** |
|  | ATOMRefine backbone | **81.69** | **70.07** | **3.81** | 54.40% |
|  | GNNRefine | 79.46 | 67.16 | 4.38 | 31.09% |
|  | ModRefiner | 81.22 | 69.09 | 4.11 | 21.76% |
| CASP14 | Initial model | 77.97 | 63.91 | 4.50 | / |
|  | ATOMRefine | **78.14** | **64.42** | 4.49 | 75.36% |
|  | ATOMRefine backbone | 78.12 | 64.40 | **4.43** | **85.51%** |
|  | GNNRefine | 77.54 | 63.60 | 4.73 | 50.72% |
|  | ModRefiner | 77.51 | 63.28 | 4.55 | 26.09% |

On average, both ATOMRefine and ATOMRefine_backbone improve the quality of backbone atoms over the initial models in terms of the GDT-TS score, GDT-HA score, and RMSD of the Cα atoms. Even though the improvement in the backbone quality is small, the results are still significant because the recent 14^th^ community-wide Critical Assessment of Techniques of Protein Structure Prediction (CASP14)^17^ showed that few refinement methods can improve the quality of the backbone of initial models on average. ATOMRefine and ATOMRefine_backbone achieve very similar performance, indicating that extending the small backbone representation to the full-atom representation for refinement still maintains the effectiveness of refining the backbones of protein structures despite that the latter needs to accommodate the extra side-chain atom refinement.

Both ATOMRefine and ATOMRefine_backbone perform better than GNNRefine and ModRefiner in terms of all three metrics on average. For instance, on the AlphaFold DB test set, the average GDT-HA scores of ATOMRefine are 0.95% higher than the following best external method ModRefiner. On the CASP14 test dataset, the average GDT-HA scores of ATOMRefine are 0.82% higher than the following best external method GNNRefine. The RMSD of ATOMRefine refined models for the AlphaFoldDB test set and CASP14 dataset is 4.08 and 4.49 Angstrom respectively, lower than 4.38 and 4.73 Angstrom of GNNRefine. The t-test shows that the difference between the initial models and ATOMRefine models in terms of the average GDT-HA score is statistically significant (P-value = 2.61E-10 on the AlphaFoldDB test set and P-value = 1.90E-08 on the CASP14 dataset). On average, out of the four methods, only ATOMRefine and ATOMRefine_backbone improve the backbone atom quality of the initial models. **Fig. 1** illustrates the change in the GDT-HA score of the refined model with respect to the initial model of these methods. ATOMRefine and ATOMRefine_backbone improve the quality of the majority of the initial models in terms of GDT-HA score.

**Fig. 1.**
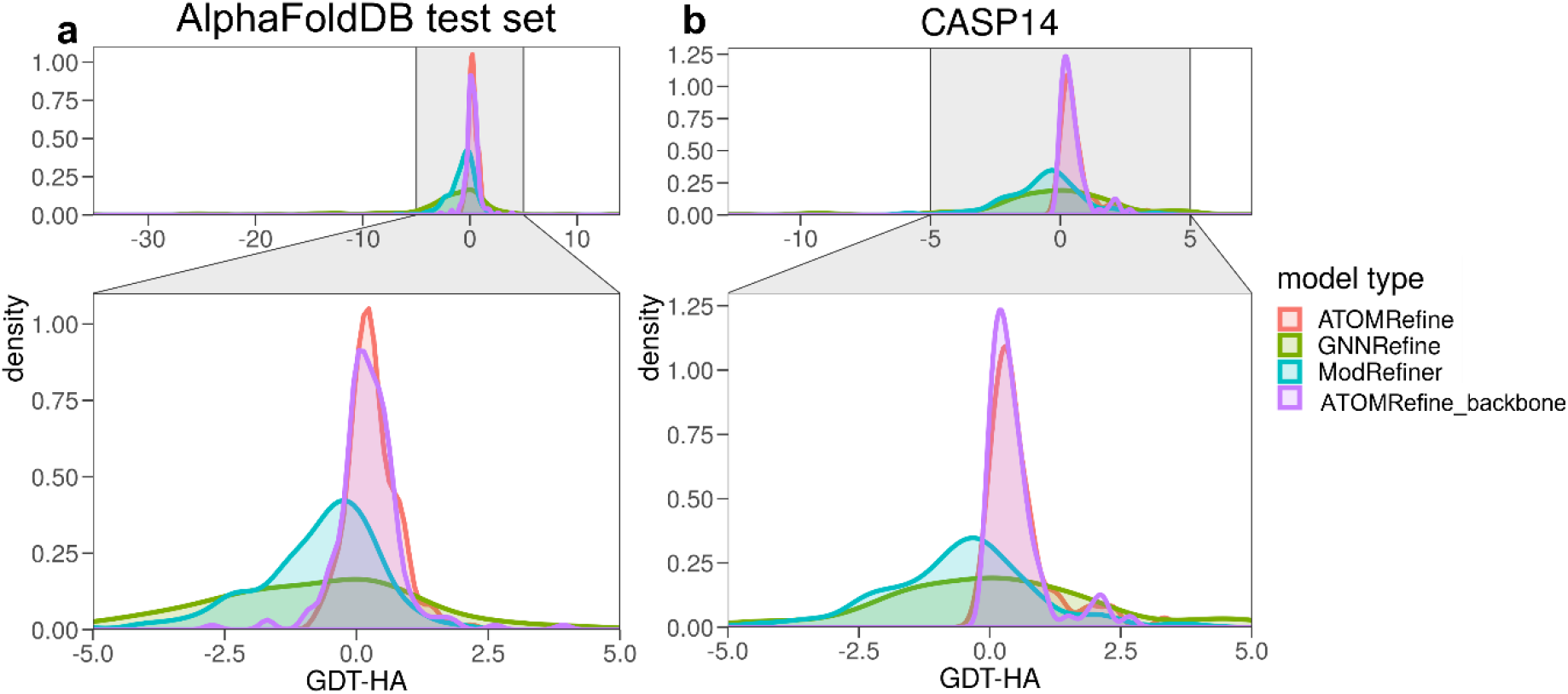
The distribution of quality change (ΔGDT-HA score) of refined models of ATOMRefine, GNNRefine, ModRefiner, and ATOMRefine_backbone with respect to the initial models. The positive value means the model quality after refinement improves from the starting model and the negative value means the model quality decreases. **a**. Results on the AlphaFold DB test set. **b**. Results on the CASP14 dataset.

### Comparison of ATOMRefine with existing methods in terms of all-atom quality

To further investigate the performance of ATOMRefine as a full-atom model refinement method, we compare ATOMRefine and ModRefiner in terms of MolProbity score based on the analysis of all-atom contacts, bond length, atom clashes, torsion angles, and side-chain rotamers. A lower MolProbity score indicates better all-atom quality and higher nativeness of the protein structure. The MolProbity score has been widely used to assess the geometric correctness and nativeness of experimentally determined protein structures before they are deposited into the PDB. To run ModRefiner, its strength parameter is set to 80 (Strength value ranges in [0,100]. Larger value makes the final model closer to the reference model). We also include GNNRefine in the full-atom level comparison. Though GNNRefine mainly focuses on refining the predicted distances of the backbone atoms, it constructs the final full-atom protein model by using the Rosetta module FastRelax^18^.

The average MolProbity scores of the initial models and the refined models of the three methods on the AlphaFold DB test set and CASP14 dataset are reported in **Fig. 2**. The average MolProbity score of the AlphaFold DB test set and CASP14 dataset is 1.31 and 1.49, much lower than 2.08 and 3.29 of the initial models, indicating a large room for improvement in the protein geometry and nativeness of the structures predicted by AlphaFold. However, the previous studies focused mostly on improving the backbone quality of protein structural models while largely ignoring enhancing their nativeness and geometry. From the results shown on **Fig. 2**, ATOMRefine also outperforms GNNRefine and ModRefiner which are also able to improve all-atom quality to some degree.

**Fig. 2.**
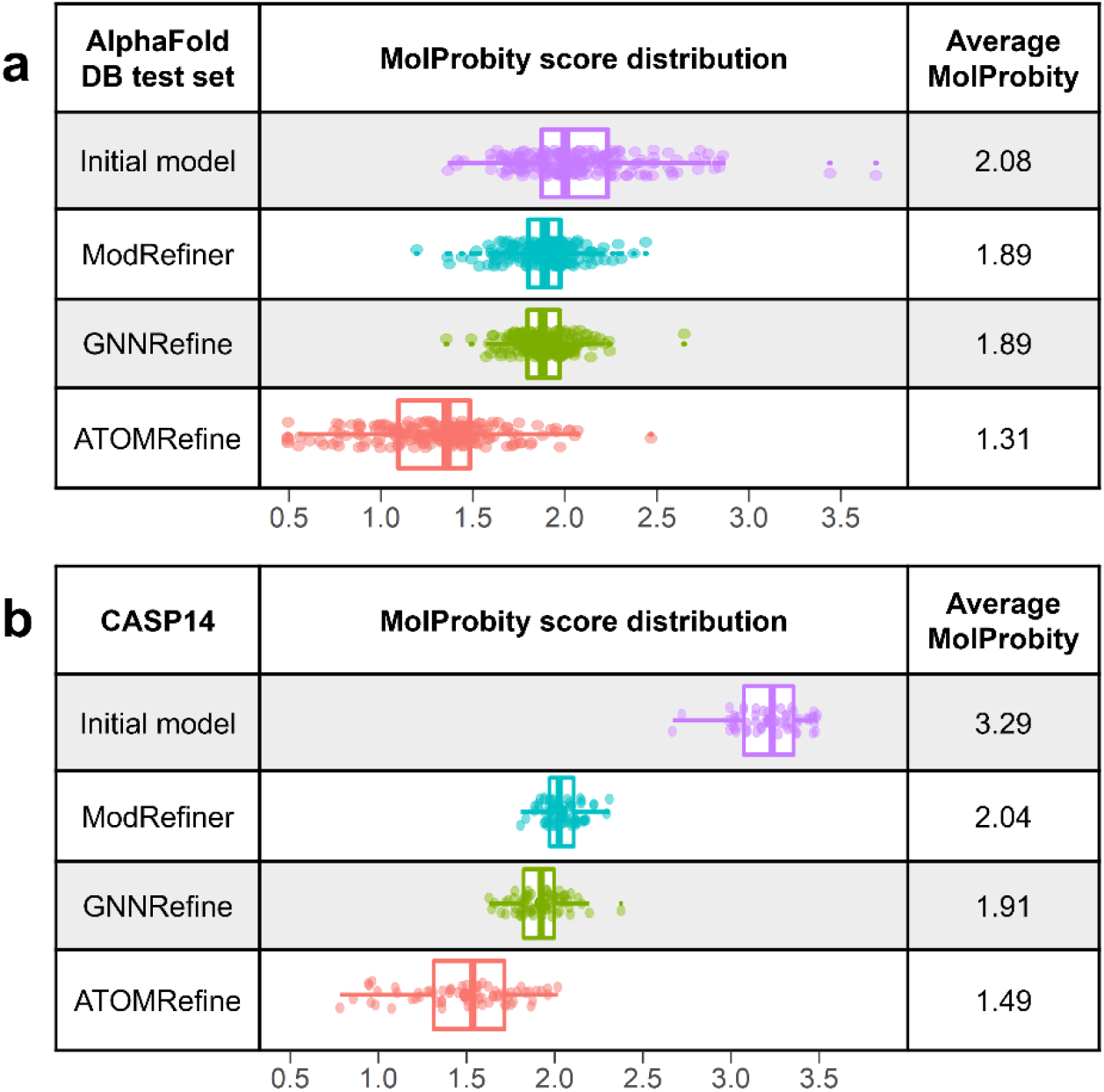
The average all-atom MolProbity score and the MolProbity score distribution of initial models and refined models of ATOMRefine, ModRefiner, and GNNRefine on **a**. AlphaFold DB test set and **b**. CASP14 dataset.

### Performance of ATOMRefine on different kinds of initial models

The outcome of model refinement is related to the quality of initial models. CASP14 official refinement targets were carefully selected by CASP organizers to assess the refinement methods considering the quality of initial models and refinement potential. In order to test the room for improvement for different targets, CASP14 selected seven targets each with an initial structure predicted by AF2 (AlphaFold2 group during CASP14 experiment) and a typical structure predicted by one of the other CASP14 groups. Therefore, each target has two different versions (v1/v2: AF2 initial model or other initial model), resulting in 14 models for refinement. In addition, those targets were classified into categories based on their modeling difficulty (FM: free modeling that does not have homologous templates in PDB, hardest targets; FM/TBM: targets in between FM and template-based modeling (TBM), second hardest; and TBM-hard: difficult TBM targets whose homologous templates exist in PDB, but are hard to find, third hardest). The name, length, classification, and initial model type can be found in **Table 2**. For each target, the GDT-HA scores of the initial models, ATOMRefine, GNNRefine, and ModRefiner are reported in **Table 2**, respectively. **Table S1** presents the results of **Table 2** according to the types of the initial models in terms of the GDT-HA score.

**Table 2.** Performance of ATOMRefine, GNNRefine, and ModRefiner on seven CASP14 refinement targets with different starting models evaluated by the GDT-HA score.

| Target ID | Residues | Classification | Initial model type | Start model | ATOMRefine | GNNRefine | ModRefiner |
| --- | --- | --- | --- | --- | --- | --- | --- |
| R1040 v1 | 130 | FM | AF2 | 54.88 | 54.62 | 50.00 | 52.12 |
| R1040 v2 |  |  | TS435_2 | 29.30 | 30.77 | 32.31 | 30.77 |
| R1041 v1 | 242 | FM | AF2 | 70.33 | 70.33 | 70.33 | 71.22 |
| R1041 v2 |  |  | TS031_1 | 40.67 | 40.89 | 40.44 | 40.22 |
| R1042 v1 | 276 | FM | TS403_1 | 34.68 | 34.88 | 34.78 | 33.87 |
| R1042 v2 |  |  | AF2 | 62.70 | 63.00 | 62.00 | 62.90 |
| R1043 v1 | 148 | FM | TS403_1 | 44.09 | 43.92 | 43.75 | 42.91 |
| R1043 v2 |  |  | AF2 | 64.86 | 65.54 | 63.34 | 64.19 |
| R1053 v1 | 171 | FM/TBM | TS042_5-D2 | 52.63 | 52.78 | 53.07 | 52.34 |
| R1053 v2 |  |  | AF2 | 79.53 | 80.26 | 73.10 | 77.05 |
| R1067 v1 | 221 | TBM-hard | TS473_3 | 45.48 | 45.14 | 45.48 | 45.14 |
| R1067 v2 |  |  | AF2 | 78.85 | 79.19 | 78.51 | 77.71 |
| R1074 v1 | 132 | FM | AF2 | 78.22 | 78.60 | 76.33 | 77.65 |
| R1074 v2 |  |  | TS140_5 | 35.42 | 35.42 | 35.42 | 33.90 |
| Average |  |  | AF2 | 69.91 | 70.22 | 67.66 | 68.98 |
|  |  |  | CASP14 other groups | 40.32 | 40.54 | 40.75 | 39.88 |

Overall, in terms of the average GDT-HA score or the GDT-HA score variation shown in **Fig. 3**, ATOMRefine outperforms GNNRefine and ModRefiner on most or all targets, respectively. With AF2 models as the initial models shown in **Fig. 3a**, the average GDT-HA score of ATOMRefine is 70.22, better than the performance of GNNRefine (67.66) and ModRefiner (68.98). ATOMRefine improves the quality of the start AF2 models whose average GDT-HA score is 69.91, but GNNRefine and ModRefiner’s GDT-HA score is lower than the GDT-HA score of the start models by 3.23% and 1.33%, respectively. With other CASP14 group models as the initial models shown in **Fig. 3b**, the average GDT-HA score of ATOMRefine is 40.54, slightly higher than 40.32 of the start models, better than 39.88 of ModRefiner, but slightly lower than GNNRefine (40.75).

**Fig. 3.**
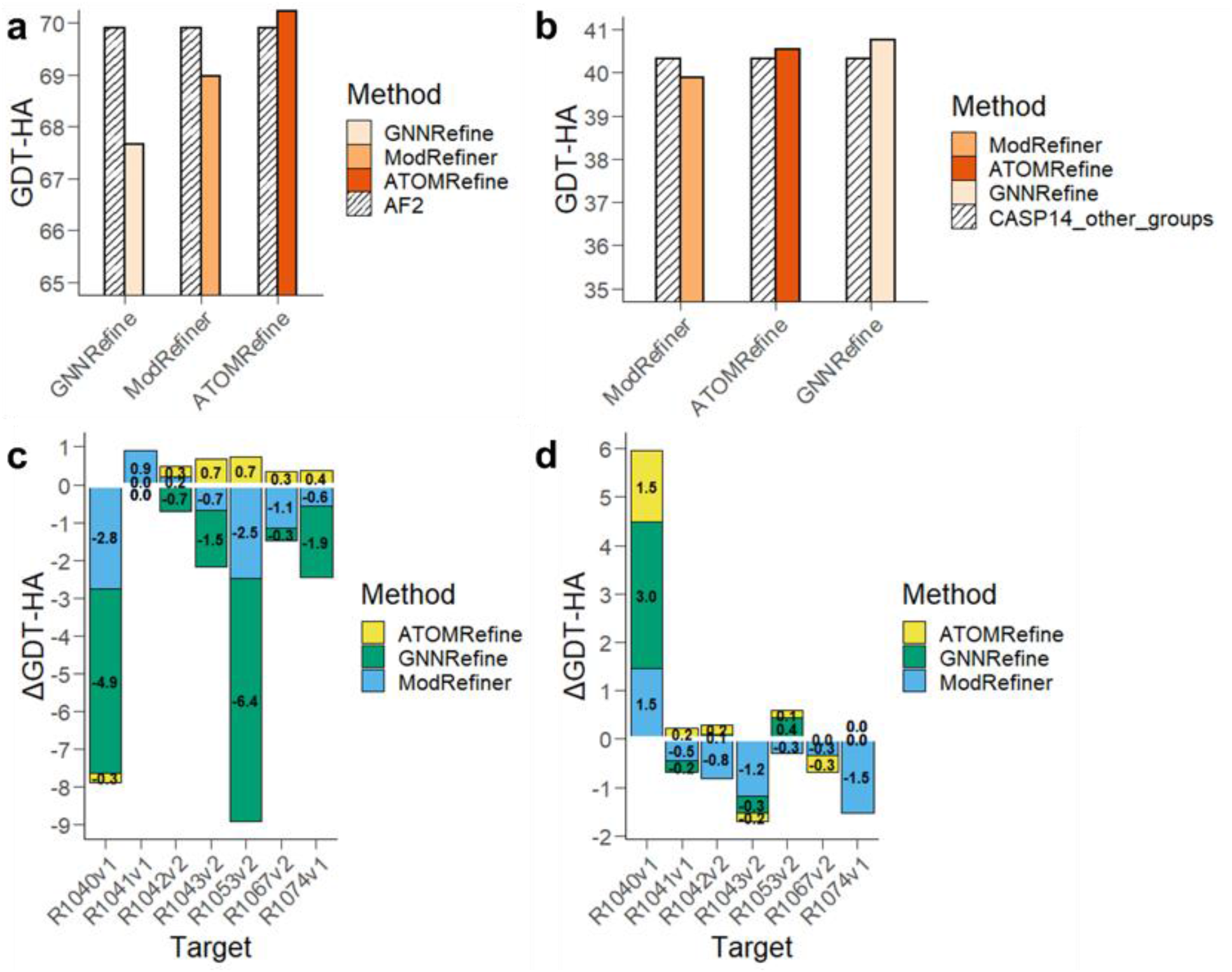
Performance of ATOMRefine, GNNRefine, and ModRefiner on seven CASP14 refinement targets by using two types of initial models as input evaluated by the GDT-HA score. **a**. The average performance of the three methods compared with the initial models of AF2. **b**. The average performance of the three methods compared with the initial models of other CASP14 groups. **c**. GDT-HA score variation of three methods with respect to the initial models of AF2. **d**. GDT-HA score variation of three methods with respective the initial models of other CASP14 groups. (GDT-HA score variation values for each method are listed on **c** and **d**; positive values stand for model improvement and negative values stand for model degradation.)

In **Fig. 3c** and **Fig. 3d**, we also list the GDT-HA score variations by applying the three refinement methods, compared to the initial models starting from either AF2 or other CASP14 groups (the specific variation values for three methods are listed in the figure). For the initial models starting from AF2, ATOMRefine produces much fewer degraded models than the other two methods. Six out of seven ATOMRefine models achieve equal or better model quality, while GNNRefine and ModRefiner show model degradation in most cases. Though GNNRefine performs better than ATOMRefine in terms of the average GDT-HA score on the initial model starting from other CASP14 groups, the number of cases achieving equal or better model quality from the two methods are the same. In general, ATOMRefine is able to maintain the model quality or improve the model quality in most cases, regardless of the types of start models.

### Comparison of the speed of ATOMRefine with other methods

In addition to maintaining or improving the model quality, ATOMRefine is also significantly faster than GNNRefine and ModRefiner. We tested the runtime of ATOMRefine, GNNRefine, and ModRefiner on the CASP14 targets with sequence length < 300. Table 3 gives the average runtime for each CASP14 target. For a protein with an average length of 156, ATOMRefine typically requires 90 seconds to complete the entire refinement process on a single Tesla V100 GPU, which is about three times faster than GNNRefine, ten times faster than ModRefiner.

**Table 3.** Runtimes on CASP14 test set

| **Table 3. Runtimes on CASP14 test set** |  |
| --- | --- |
| **Method** | **Total runtime(s)** |
| ATOMRefine | 90 |
| GNNRefine | 335 |
| ModRefiner | 974 |

## Conclusion

In this paper, we introduce ATOMRefine, a novel full-atom 3D-equivariant graph transformer method for protein structure refinement. It uses a new full-atom graph to represent atoms, bonds, and coordinates as the node and edge features, which is processed by the equivariant and invariant layers of the SE(3) graph transformer to refine the coordinates of all the atoms. We rigorously evaluate ATOMRefine on three test datasets. Compared to the refinement methods focusing on refining backbone atoms, it has the advantage of directly generating an all-atom refined structure. Moreover, ATOMRefines can improve the quality of both backbones and all atoms including side-chain atoms over the initial input models and outperforms the state-of-the-art deep learning and energy minimization-based methods. Finally, once it is trained, ATOMRefine can refine protein structure very quickly, making it applicable to proteome-wide protein structure refinement.

We plan to further improve ATOMRefine by training it on a larger dataset consisting of AlphaFold models of more diverse quality, particularly including more low-quality models. In the current training dataset, 92% of structural models are high-accuracy models, which may limit the amount of improvement that can be made by the deep learning method. Adding more low-quality models into training may make ATOMRefine learn to make larger improvements to the backbone structure on less accurate input.

## Methods

ATOMRefine is an end-to-end protein refinement method based on a SE (3)-equivariant graph neural network. It directly predicts refined atomic coordinates of all the atoms as output from the initial coordinates of all the atoms in an input structure. To avoid the bond geometry violation, a final relaxation step by Amber^19^ is added in the ATOMRefine pipeline. Its simplified version based on the same deep learning architecture is used to refine the coordinates of backbone atoms only. The overall framework of ATOMRefine is illustrated in **Fig. 4**. Details of the graph representation, network architecture, training and test data, and evaluation metrics are described as follows.

**Fig. 4.**
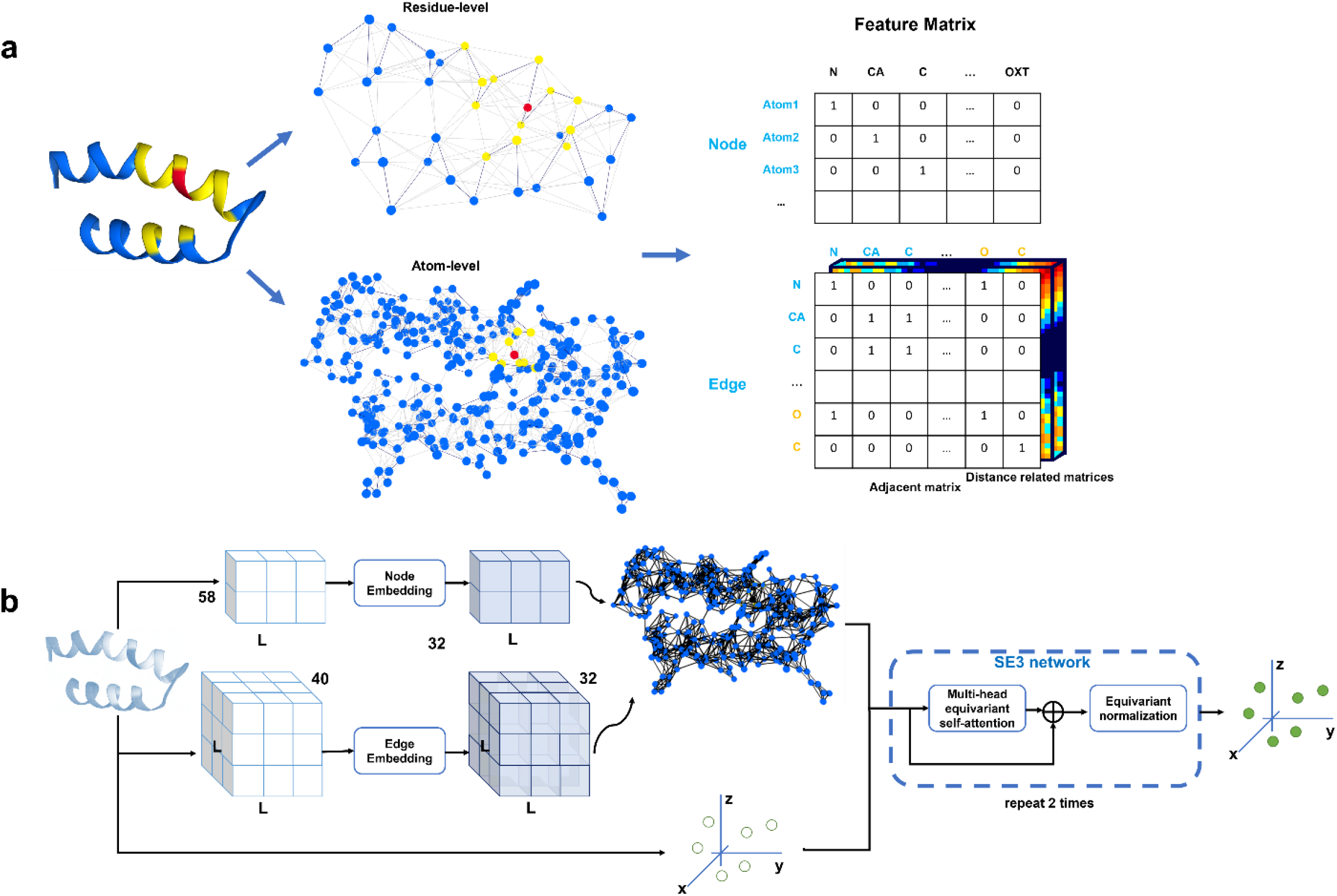
The ATOMRefine framework. **a**. ATOMRefine graph representation of a protein structure at all-atom level or backbone atom-level. A node is used to represent an atom. The 3D protein structure is encoded as the atomic features (node features) and inter-atom features (edge features including adjacent bond matrix and distance-related matrices). Each node (e.g., a node in red) is connected to the *k* nearest neighboring nodes (nodes in yellow) selected by the Euclidean distance calculated from atom 3D coordinates. The covalent bond edge between atoms shown as the solid blue line in the graph is also an edge feature stored in the adjacent bond matrix. **b**. the deep learning architecture of ATOMRefine. Each block in the SE(3) transformer network consists of one equivariant GCN attention block and one SE(3)-equivariant normalization layer.

### All-atom graph-based representation of protein structure and SE(3) graph transformer architecture

In this work, a protein structure is considered as a set of nodes each of which represents an atom in the protein. Each atom *i* has a 3D coordinate (x_*i*_, y_*i*_, z_*i*_) that can be used to calculate the pairwise spatial relations between atoms. A protein structure is represented as a graph of the nodes in which the edges describe the relationships between the nodes (i.e., atoms).

Each node has atom features including one-hot encoding of atom types (a binary vector indicating 37 atom types) and the types of amino acids that the atom belongs to. Each node also has x, y, z coordinates as variable features that will be updated.

As illustrated in Fig. 4a, each node is connected to the *k* (*k* = 128) nearest neighboring nodes selected by the Euclidean distance between atom 3D coordinates through edges.

Six edge features are generated, including one distance-based edge feature, one covalent bond edge feature, one relative position edge feature, and three relative orientation edge features. For the distance-based edge feature, we use the radial basis function to convert the distance (*d*) between two nodes as features: 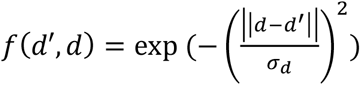, where *d* is the Euclidean distance between two nodes of an edge, d’ and *σ*_*d*_ are hyperparameters. We set *σ*_*d*_ = 0.57, d’ = [0, *σ*_*d*_, 2*σ*_*d*_, …, 35*σ*_*d*_] ∈ [0Å, 20Å], following the work of RoseTTAFold. So, for each edge, there are 36 distance-based edges.

We the covalent bond edge feature to represent the local covalent bond connectivity between atoms. An adjacent bond matrix (M) is calculated from the atom-atom distance matrix (D) to detect if there is a covalent bond between two atoms according to the work of Graphein^20^. We parse the atomic Euclidean distance matrix ***D*** to the binary covalent bond adjacent matrix ***M*** as shown in Equation 1,

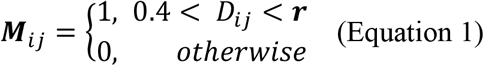

where i, j are the atom positions (indices), and the thresholding parameter *r* is a set of covalent radii based on different atom types. 1 indicates there is a bond between two atoms.

Similar to the work of *Octavian-Eugen Ganea*^21^ and *trRosetta*^22^, we also use the relative position and relative orientation features for edges based on the local coordinate system. We construct the local coordinate system based on each amino acid residue position (index) in a protein model (for atoms of the same amino acid, they share the same local coordinate basis). As shown in **Fig. 5**, for each residue *i*, we define the Cα coordinate as the origin, the unit vector pointing from Cα atom to C atom as *u*_*i*_, and the unit vector pointing from Cα atom to N atom as *y*_*i*_ (on the y axis). The normal of the plane C-Cα-N is defined as *z*_i_ (on the z-axis), where 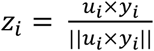. Naturally, we define *x*_*i*_ = *y*_*i*_ × *z*_*i*_ (on the x-axis). In total, *x*_*i*_, *y*_*i*_ and *z*_*i*_ consist of the basis of residue *i*’s local coordinate system. As shown in Equation 2, the relative position edge feature *p*_*im,jn*_ denotes the relative position of atom *n* in the residue position *j* to atom *m* in the residue *i*. ***atom***_*jn*_ denotes the coordinate of atom *j* in the residue position *n*.

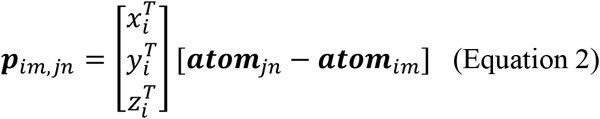

**Fig. 5.**
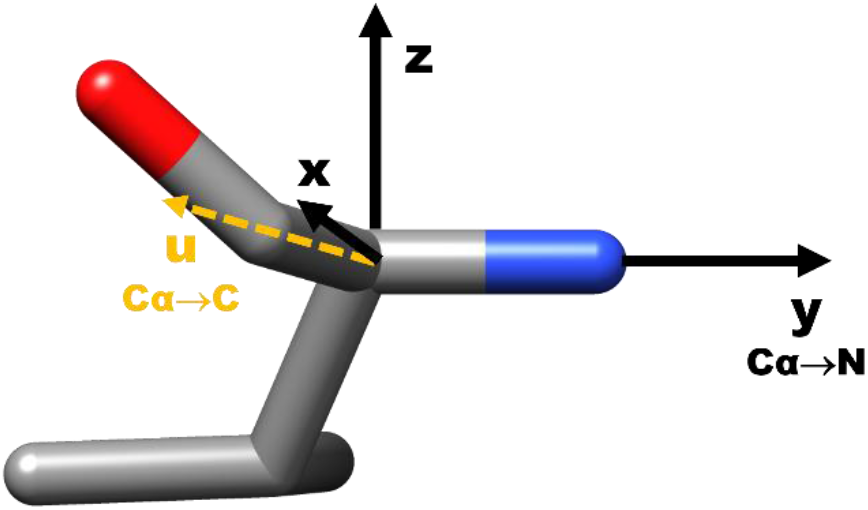
Representation of a single valine residue and its local coordinate system. We define the atom Cα as the origin, The y-axis points from the atom Cα to the atom N. The x-axis is placed in the plane of C-Cα-N. Following the right-hand coordinate system, the z-axis is the normal of the plane.

As shown in Equation 3, relative orientation features *q*_*im,jn*_, *k*_*im,jn*_, *t*_*im,jn*_ denote the relative orientation of atom *n* in residue position *j* to atom *m* in the residue position *i*.

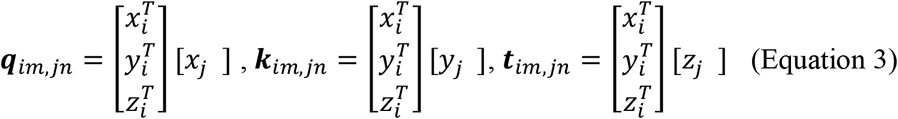

With atom and atom-atom relationship features encoded as node features and edge features above, the protein structure can be encoded in the graph. **Fig. 4a** shows the scheme of the graph representation of a protein model at the atom level. The detailed atomic and residue-based features are presented in **Table S2**.

The general network architecture of ATOMRefine is illustrated in **Fig. 4b**. We parse the initial protein model as the node and edge features to build a graph representation. The graph is then fed into the SE(3)-transformer to refine the given 3D atom coordinates. All features of each node except for 3D coordinates correspond to SE (3) type 0 node feature, and the 3D coordinates of each node (atom) correspond to SE (3) type 1 node feature. The embedding size of the input node and edge features are set to 32. A SE (3) transformer is used to predict the coordinate shifts between the initial model and the native structure. The SE(3) transformer consists of two SE3 equivariant attention blocks, including one multi-head attention block with 16 channels and 4 attention heads, and one SE3 equivariant normalization layer. For each attention block, queries are the linear projection of the graph node features. Keys and values are from graph edge features computed by TFN^23^ layers. ATOMRefine is implemented on top of the deep learning framework PyTorch Lightning^24^ and Deep Graph Library^25^.

#### Training data

We download the predicted protein models from AlphaFold DB. We use MMseqs2^26^ to remove the sequence duplication first and then match the remaining protein sequences with the native structures that exist in Protein Data Bank. A structural model is matched with a true structure if the following criteria are met: (a) the model sequence matches with the native sequence; (b) protein sequence length >= 50. In total, 13,121 are selected as the initial models and their true structures are used as labels. ATOMREfine is trained and validated on the training data via 10-fold cross-validation. During training, protein structures with >1500 residues are cropped to fit the GPU memory. We set Adam as the optimizer with parameters: β1=0.9, β2=0.999, and weight decay=0.001. We set the batch size as 1 and use the mean squared error between predicted coordinates and true coordinates of atoms as the loss function. We set the number of training epochs to 50 with the early stopping when there are no improvements in the validation loss for five consecutive epochs. Ten ATOMRefine models have been generated accordingly.

#### Test data

We use three test datasets to evaluate the methods: an AlphaFold DB test set containing 193 protein targets retrieved from the Alphafold DB, the CASP14 dataset containing 69 regular targets, and the CASP14 refinement dataset (7 protein targets). The CASP14 refinement targets are a subset of the CASP14 regular targets, selected by CASP organizers for challenging predictors to make a structural refinement. Any sequences in the training data that has >=30% identity with any sequences in the three test datasets have been removed in the training data preparation so that there is no overlap between the training data and the three all three test sets (e.g., sequence identity < 30%). For each target in the CASP14 dataset, Alphafold2 is run to generate start models. For the CASP14 refinement dataset, we use the initial models provided by CASP14 organizers as the star models. All the true structures for the targets in the test datasets are obtained from the PDB.

### Evaluation metrics

To compare the model quality of initial models and refined models, we use GDT-HA^27^, GDT-TS^27^, RMSD of the Cα atoms, and Molprobity score^28^ as four main evaluation metrics. GDT-TS is the global distance score. It ranges from 0 to 100% (or simply from 0 to 1), a higher value indicating better model accuracy. GDT-HA is the high–accuracy version of the GDT-TS score with smaller distance cutoffs. RMSD of the Cα atoms measures the root mean square deviation of the Cα atoms in a protein model from its native structure, describing the accuracy of the positions of the Cα atoms. A lower RMSD means better quality. The MolProbity score assesses the quality of all the atoms of a model including side-chain atoms. It considers atom contacts, atom clashes, bond lengths and angles, and torsion angles. A lower Molprobity score indicates better model quality.

## References

1. Jumper, J. et al. Highly accurate protein structure prediction with AlphaFold. Nature 596, 583–589 (2021).

2. Senior, A. W. et al. Improved protein structure prediction using potentials from deep learning. Nature 577, 706–710 (2020).

3. Tunyasuvunakool, K. et al. Highly accurate protein structure prediction for the human proteome. Nature 596, 590–596 (2021).

4. Cretin, G., Galochkina, T., de Brevern, A. G. & Gelly, J.-C. PYTHIA: Deep Learning Approach for Local Protein Conformation Prediction. Int J Mol Sci 22, 8831 (2021).

5. Jones, D. T. & Thornton, J. M. The impact of AlphaFold2 one year on. Nature Methods 19, 15–20 (2022).

6. Pearce, R. & Zhang, Y. Toward the solution of the protein structure prediction problem. Journal of Biological Chemistry 297, 100870 (2021).

7. Bhattacharya, D. & Cheng, J. 3Drefine: Consistent protein structure refinement by optimizing hydrogen bonding network and atomic-level energy minimization. Proteins: Structure, Function, and Bioinformatics 81, 119–131 (2013).

8. Heo, L., Janson, G. & Feig, M. Physics-based protein structure refinement in the era of artificial intelligence. Proteins: Structure, Function, and Bioinformatics 89, 1870–1887 (2021).

9. Mirjalili, V., Noyes, K. & Feig, M. Physics-based protein structure refinement through multiple molecular dynamics trajectories and structure averaging. Proteins: Structure, Function, and Bioinformatics 82, 196–207 (2014).

10. Heo, L., Park, H. & Seok, C. GalaxyRefine: Protein structure refinement driven by side-chain repacking. Nucleic Acids Res 41, W384–W388 (2013).

11. Lee, G. R., Won, J., Heo, L. & Seok, C. GalaxyRefine2: simultaneous refinement of inaccurate local regions and overall protein structure. Nucleic Acids Research 47, W451– W455 (2019).

12. Xu, D. & Zhang, Y. Improving the physical realism and structural accuracy of protein models by a two-step atomic-level energy minimization. Biophys J 101, 2525–2534 (2011).

13. Hiranuma, N. et al. Improved protein structure refinement guided by deep learning based accuracy estimation. Nature Communications 12, 1340 (2021).

14. Jing, X. & Xu, J. Fast and effective protein model refinement using deep graph neural networks. Nature Computational Science 1, 462–469 (2021).

15. Minkyung, B. et al. Accurate prediction of protein structures and interactions using a three-track neural network. Science (1979) 373, 871–876 (2021).

16. Fuchs, F., Worrall, D., Fischer, V. & Welling, M. Se (3)-transformers: 3d roto-translation equivariant attention networks. Advances in Neural Information Processing Systems 33, 1970–1981 (2020).

17. Simpkin, A. J., Sánchez Rodríguez, F., Mesdaghi, S., Kryshtafovych, A. & Rigden, D. J. Evaluation of model refinement in CASP14. Proteins: Structure, Function, and Bioinformatics 89, 1852–1869 (2021).

18. Chaudhury, S., Lyskov, S. & Gray, J. J. PyRosetta: a script-based interface for implementing molecular modeling algorithms using Rosetta. Bioinformatics 26, 689–691 (2010).

19. Salomon-Ferrer, R., Case, D. A. & Walker, R. C. An overview of the Amber biomolecular simulation package. WIREs Computational Molecular Science 3, 198–210 (2013).

20. Jamasb, A. R. et al. Graphein - a Python Library for Geometric Deep Learning and Network Analysis on Protein Structures and Interaction Networks. bioRxiv 2020.07.15.204701 (2021) doi:10.1101/2020.07.15.204701.

21. Ganea, O.-E. et al. Independent SE (3)-Equivariant Models for End-to-End Rigid Protein Docking. arXiv preprint arXiv:2111.07786 (2021).

22. Jianyi, Y. et al. Improved protein structure prediction using predicted interresidue orientations. Proceedings of the National Academy of Sciences 117, 1496–1503 (2020).

23. Thomas, N. et al. Tensor field networks: Rotation-and translation-equivariant neural networks for 3d point clouds. arXiv preprint arXiv:1802.08219 (2018).

24. Falcon, W. & The PyTorch Lightning team. PyTorch Lightning. Preprint at https://doi.org/10.5281/zenodo.3828935 (2019).

25. Wang, M. et al. Deep graph library: A graph-centric, highly-performant package for graph neural networks. arXiv preprint arXiv:1909.01315 (2019).

26. Steinegger, M. & Söding, J. MMseqs2 enables sensitive protein sequence searching for the analysis of massive data sets. Nature Biotechnology 35, 1026–1028 (2017).

27. Zemla, A. LGA: a method for finding 3D similarities in protein structures. Nucleic Acids Research 31, 3370–3374 (2003).

28. Williams, C. J. et al. MolProbity: More and better reference data for improved all-atom structure validation. Protein Science 27, 293–315 (2018).

