## Supplemental Table S1 and S2 for "Atomic protein structure refinement using all-atom graph representations and SE(3)-equivariant graph neural networks"

| **Table S1 Performance of ATOMRefine, GNNRefine, and ModRefiner on seven CASP14 refinement targets with different starting models. (a)** AF2 start model and evaluated by GDT-HA score; **(b)** other CASP14 start model and evaluated by GDT-HA score; | | | | | | |
| --- | --- | --- | --- | --- | --- | --- |
| **a. AF2 start model and evaluated by the GDT-HA score** | | | | | | |
| **TargetID** | **Residues**  **in**  **refinement target** | **Classification** | **Starting model**  GDT-HA | **ATOMRefine** GDT-HA | **GNNRefine** GDT-HA | **ModRefiner**  GDT-HA |
| R1040v1 | 130 | FM | **54.88** | 54.62 | 50.00 | 52.12 |
| R1041v1 | 242 | FM | 70.33 | 70.33 | 70.33 | **71.22** |
| R1042v2 | 276 | FM | 62.70 | **63.00** | 62.00 | 62.9 |
| R1043v2 | 148 | FM | 64.86 | **65.54** | 63.34 | 64.19 |
| R1053v2 | 171 | FM/TBM | 79.53 | **80.26** | 73.10 | 77.05 |
| R1067v2 | 221 | TBM-hard | 78.85 | **79.19** | 78.51 | 77.71 |
| R1074v1 | 132 | FM | 78.22 | **78.6** | 76.33 | 77.65 |
| **b. Other CASP14 group start model and evaluated by the GDT-HA score** | | | | | | |
| **TargetID** | **Residues**  **in**  **refinement target** | **Classification** | **Starting model**  GDT-HA | **ATOMRefine** GDT-HA | **GNNRefine** GDT-HA | **ModRefiner**  GDT-HA |
| R1040v2 | 130 | FM | 29.3 | 30.77 | **32.31** | 30.77 |
| R1041v2 | 242 | FM | 40.67 | **40.89** | 40.44 | 40.22 |
| R1042v1 | 276 | FM | 34.68 | **34.88** | 34.78 | 33.87 |
| R1043v1 | 148 | FM | **44.09** | 43.92 | 43.75 | 42.91 |
| R1053v1 | 171 | FM/TBM | 52.63 | 52.78 | **53.07** | 52.34 |
| R1067v1 | 221 | TBM-hard | **45.48** | 45.14 | **45.48** | 45.14 |
| R1074v2 | 132 | FM | **35.42** | **35.42** | **35.42** | 33.9 |

| **Table S2 List of all atom-level and backbone atom-level features.** | | | |
| --- | --- | --- | --- |
| **Methods** | **Feature Type** | **Feature Name** | **Details** |
| All atom-level features  for ATOMRefine | Node feature | Atom Embedding | 37 atom types (N, CA, C, O, CB, OG, CG, CD1, CD2, CE1, CE2, CZ, OD1, ND2, CG1, CG2, CD, CE, NZ, OD2, OE1, NE2, OE2, OH, NE, NH1, NH2, OG1, SD, ND1, SG, NE1, CE3, CZ2, CZ3, CH2, OXT) + 21 AA types (20 natural amino acids and one for unknown) |
|  | Edge feature | Distance-Based Edge Features | Inter-atom distance matrices |
|  |  | Covalent bond matrix | The distance matrix is then thresholded to entries less than this distance plus some tolerance to create and adjacency bond matrix. This adjacency bond matrix is then parsed into an edge list. |
|  |  | Relative Edge Features | Relative Position Edge Features  Relative Orientation Edge Features |
| Backbone atom-level (or residue level) features for ATOMRefine_backbone | Node feature | Amino Acid Embedding | One-hot encoding of 21 AA types (20 natural amino acids and one for unknown) |
|  |  | Dihedral angle | In radian |
|  |  | Secondary structure | 3 states |
|  |  | Relative solvent accessibility | Calculated by DSSP and ranges from [0,1] |
|  | Edge feature | Distance-Based Edge Features | Distance maps (Cα - Cα, Cβ - Cβ and N - O) |
|  |  | Orientation features map | ω, θ, φ calculated by trRosetta |

Note: the x, y, z coordinate of atom is also a node feature that is updated by the network, which is not shown in the table above.
